# Learning supervised embeddings for large scale sequence comparisons

**DOI:** 10.1101/620153

**Authors:** Dhananjay Kimothi, Pravesh Biyani, James M Hogan, Akshay Soni, Wayne Kelly

## Abstract

Similarity-based search of sequence collections is a core task in bioinformatics, one dominated for most of the genomic era by exact and heuristic alignment-based algorithms. However, even efficient heuristics such as BLAST may not scale to the data sets now emerging, motivating a range of alignment-free alternatives exploiting the underlying lexical structure of each sequence.

In this paper, we introduce `SuperVec`, a novel supervised approach to learning sequence embeddings. Our method extends earlier Representation Learning (RL) based methods to include jointly contextual and class-related information for each sequence during training. This ensures that related sequence fragments have proximal representations in the target space, better reflecting the structure of the domain.

Such representations may be used for downstream machine learning tasks or employed directly. Here, we apply `SuperVec` embeddings to a sequence retrieval task, where the goal is to retrieve sequences with the same family label as a given query. The `SuperVec` approach is extended further through `H-SuperVec`, a tree-based hierarchical method which learns embeddings across a range of feature spaces based on the class labels and their exclusive and exhaustive subsets.

Experiments show that supervised learning of embeddings based on sequence labels using `SuperVec` and `H-SuperVec` provides a substantial improvement in retrieval performance over existing (unsupervised) RL-based approaches. Further, the new methods are an order of magnitude faster than BLAST for the database retrieval task, supporting hybrid approaches in which `SuperVec` rapidly filters the collection so that only potentially relevant records remain, allowing slower, more accurate methods to be executed quickly over a far smaller dataset. Thus, we may achieve faster query processing and higher precision than before.

Finally, for some problems, direct use of embeddings is already sufficient to yield high levels of precision and recall. Extending this work to encompass weaker homology is the subject of ongoing research.

## 1 Introduction

Rapid comparison of molecular sequences is an essential task in bioinformatics, with applications including homology detection, annotation, and phylogenetic analysis [1]. For most of the genomic era, sequence comparison has relied primarily on a succession of exact and subsequently heuristic algorithms for sequence alignment, the most successful being BLAST, the *Basic Local Alignment Search Tool*, which has in various guises dominated the field since its introduction in 1990 [2]. Exact algorithms for finding global or local sequence alignments were introduced respectively in [3] and [4]. Here the alignment score constitutes a pseudo-metric and a principled, yet time-consuming, method for quantifying sequence similarity and divergence. A high alignment score is taken to indicate a high probability that the sequences are similar. However, this intuition may break down for isolated local alignments or more remote homology, and fail altogether under structural re-arrangement.

BLAST rapidly identifies short, high-scoring seed matches between the sequences, extending and combining these as long as a strong match score is maintained. As is well known, the approximate alignment and its significance are estimated using Karlin-Altschul statistics, and these are used to rank the results returned when queries are submitted to sequence collections. BLAST uses word-based heuristics to avoid the computational penalties inherent in the dynamic programming based exact alignment algorithms, which are quadratic in the sequence length. Yet the exponential increase in the availability of sequence data as a result of successive ‘next-generation sequencing’ (NGS) technologies has highlighted the limitations even of successful heuristics of this kind, motivating research into comparison methods that do not rely on the underlying primitive of sequence alignment. Such alignment-free methods construct similarity through alternative features, usually based upon the bag-of-words (BOW) representation familiar from information retrieval, and have the additional virtue of robustness in the presence of structural re-arrangements.

The construction of BOW for sequences follows two steps: (i) splitting the biological sequence into *k-mers* (or words), and (ii) computing the statistics for either an individual *k-mer* or a *k-mer* set. The limitations of the BOW representation lie in its high dimensionality and inability to capture patterns inherent in the sequence, patterns which might otherwise support more meaningful comparison. Recent advances in learning distributed representations for text processing have led to techniques that provide a semantically meaningful, dense vector representation of words present in a large corpus of documents. Such Representation learning (RL) approaches in text processing rely on the distributional hypothesis of semantics, the idea that words which frequently co-occur support a common meaning. The best known of these approaches is `Word2Vec`, due to Mikolov et.al [5]. These techniques may also be applied to biological sequences to analyse, compare and perform downstream machine learning tasks for various applications. Initial research by Asgari et al. [6] and Kimothi et al. [7] utilized `Word2Vec` and `Doc2Vec` respectively to demonstrate that useful low-dimensional and robust representations can be generated to support machine learning tasks over biological sequences.

### 1.1 Contribution

This paper concerns learning of tailored word-based embeddings for molecular sequences to support sequence comparison without the need for sequence alignment. As with other embedding methods, our technique also requires as input a set of words or *k-mers* extracted from the sequence. The effect of learning is to capture the information implicit in these *k-mers*, markedly reducing the dimension from the full bag-of-words representation while ensuring that related *k-mer* groups have proximal representations in the embedding space. We improve these associations by the addition of class information through supervision, and subsequently through the use of supervision over defined class hierarchies. In this way, we may compute sequence similarity accurately over vectors within a lower dimensional subspace, while ensuring that the calculation relies on features pertinent to the problem at hand, here reflecting some biological grouping or functional relationship.

We demonstrate the utility of our proposed approach for retrieval of *homologous* sequences—sequences with a common evolutionary history—from a database. Here we take membership of the same protein family as a marker of homology, relying on the family definition provided in PFAM [8] for protein sequences. The exponential increase in database size over the last decade or more has posed fundamental challenges for alignment-based techniques, primarily due to the quadratic complexity of the algorithms. In this paper, we propose alternative techniques for homologous sequence retrieval which provide an order of magnitude improvement in execution time over these methods. The methods, which we term `SuperVec`, and in its hierarchical form, `Hierarchical SuperVec`, fall broadly under the umbrella of alignment-free sequence comparison but are based more firmly in the tradition of Representation Learning (RL) in text processing.

In the earlier bioinformatics applications ([6], [7]), *k-mers* are embedded in a manner that reinforces their surrounding context, implicitly capturing the ‘semantics’ of these co-occurring terms. Here we implicitly adopt a distributional hypothesis for sequence *k-mers*, the view that commonly occurring *k-mers* are likely drawn from functionally related contexts, as part of some region of a gene or protein, or as a constituent of some regulatory region. The nature of these relationships depends crucially on the type of the sequence and on the resolution implied by the choice of *k-mer* size, *k*. The addition of metadata to the training process both reinforces and complements the relationships inherent in the sequence representation. Such labels are commonly used to indicate function (as in the *Gene Ontology* categories [9]) or associations and may be used directly (see for example, the protein-protein interaction work of [10]) or indirectly to support inference. Class information of this nature may be available implicitly in features or representations exploited by other methods, for example in the BLOSUM scores employed by BLAST [11], which summarise the alignments of hundreds of proteins.

`SuperVec` makes supervised learning of sequence representations based on class information explicit, extending the earlier context-based ([6], [7]) approaches to incorporate additional labels or metadata associated with the sequences from which we derive a set of *k-mers*. Our hypothesis is that by infusing meta information in the learning process, the approach will yield sequence representation better suited to the task at hand, with members of the same class (even if they are divergent sequences – those who might otherwise exhibit a lower degree of shared context) – embedded in close vicinity in the vector space. This is achieved by joining two embedding models in one framework. The first enforces class supervision, while the second incorporates contextual information present in the sequence, achieved as before by extracting a set of *k-mers* from each sequence. Class information constrains the intra-class vectors to fall closer together within the vector space, which in turn induces class information in the embedding of the constituent *k-mer* sets.

The number of constraints enforced increases with the number of classes, reducing the efficacy of the training process and ultimately limiting its accuracy as the inter-class separation decreases. To overcome these issues we consider a range of partitions other than the original classes, and construct a series of embeddings using the `SuperVec` algorithm to better cover the space. This method, which we call `Hierarchical SuperVec` or `H-SuperVec`, supports embeddings across feature spaces based on the class labels and their exclusive and exhaustive subsets. As we have seen, some of the most commonly used label sets such as GO [12] are inherently hierarchical in nature, supporting the application of these partitions across multiple levels. This approach has the effect of providing observations of the separation of the underlying sequences, as projected into the particular embedding space. These may be exploited to improve our estimate of the true separation, and to better capture the structure of the original sequence hierarchy through the use of label sets reflecting these relationships. In the bioinformatics context, we may work with a number of additional label sources such as the functional categories of the Gene Ontology [12] and regulatory pathway information from databases such as KEGG [13] and RegPrecise [14].

Experimental results on sequence retrieval tasks illustrate that `SuperVec` provides a substantial improvement (40 – 100% for many precision-recall values) in the retrieval performance vis-a-vis the embedding methods reported in [6] and [7]. These results also demonstrate that `H-SuperVec` further improves the retrieval performance providing 80 – 200% improvement for many precision-recall values when compared to [6] and [7].

The proposed methods can also be used as a filter to rapidly select a relevant subset of sequences from the large database, allowing methods with high precision to give desirable output. We call such an approach as a Hybrid approach. The experimental results show that the Hybrid approach-`H-SuperVec+BLAST` is significantly faster than the BLAST and gives similar performance for early recall levels. While the representations provided by our model are used in this paper only for the homologous sequence retrieval problem, these representations can be directly utilised for other bioinformatics applications, such as the prediction of protein-protein interactions. Further, there is also some scope for improvement in learning models that provide vector space representations for biological sequences, and this paradigm has the potential to match the accuracy of alignment based methods while offering far greater computational efficiency, in part through the reduced dimensionality of the representation.

In summary, following are the main contributions of the paper:

- We present a supervised approach - `SuperVec` to learn embeddings for biological sequences. `SuperVec` provides flexibility to utilize meta-information (like class labels) along with the contextual information present in the sequences to generate their embeddings.
- We present an approach - `H-SuperVec` that is designed specifically for sequence retrieval task. `H-SuperVec` is built over series of `SuperVec` models. Although we have shown the use of `SuperVec` and `H-SuperVec` approaches for particular bioinformatics application these approaches are generic and can be easily extended to other domains with similar problems.
- We show that our approaches provide a faster alternative to the alignment-based method like BLAST for sequence retrieval task, providing an order of magnitude speedup in querying time.
- We show that our approaches can be used as a pre-processing filter for applying a high precision method for sequence retrieval tasks.

In the next section we give a more detailed introduction to the key approaches in Representation Learning, `Word2Vec` and `Doc2Vec`, providing the building blocks for the development of `SuperVec` and `H-SuperVec` in the subsequent sections.

## 2 Embedding Models for Text Documents

We have earlier introduced the general ideas of Representation Learning, their origins in text processing, and the application of these methods in a biological context. In this section we shall consider the assumptions underpinning these approaches and some of the insights gained, leading into the development of `SuperVec` and its hierarchical variant `H-SuperVec`. To assist this process, we conclude this section with a brief introduction to the architecture of `Word2Vec` and `Doc2Vec`, which have been prominent in previous studies and appear later as building blocks for our proposed methods.

The fundamental principle of `Word2Vec` [5] lies in the distributional hypothesis [15]: co-occurring words also share a semantic relationship. `Word2Vec` captures co-occurrence information in word embeddings by employing a simple classification task wherein a word is predicted based on its context (nearby words) and the representations are learned in a manner such that words with similar meanings appear proximal in the embedding space. Follow up works [16–18] extended the idea of word embedding to learning of embeddings for the whole document, based on suitable combinations of word embeddings.

Asgari et. al. [6] introduced `BioVec`, an adaptation of the `Word2Vec` framework to learn embeddings for molecular sequences. Here, the skip-gram variant of `Word2Vec` is used to obtain *k-mer* embeddings, with the sequence embedding given by the sum of the embeddings of all the *k-mers* present in the sequence. However, `BioVec` does not preserve the *k-mer* ordering of the sequence. Kimothi et.al. [7] addressed this limitation through `Seq2Vec` [7], which relies on the `Doc2Vec` [16] architecture to provide a direct embedding for the sequence.

While the application of `Word2Vec` and `Doc2Vec` to biological sequences is rather straightforward, the insights obtained are not. Both of these studies confirmed the utility of the embeddings in capturing relevant features of the protein sequences — initially using `BioVec` and through subsequent improvements in `Seq2Vec`. The `BioVec` study also provided an important insight regarding the relationship among *k-mer* embeddings and the biochemical and biophysical properties of the sequence, the mass, volume, and charge. It was shown that *k-mers* with similar physico-chemical properties form clusters in the embedding space. Further, in the `Seq2Vec` paper we demonstrated that the use of the sequence tag during learning results in even better clustering of the sequence data - sequences belonging to the same family were more likely to be clustered together using `Seq2Vec` than with the `BioVec` model. Such representations have proven beneficial for classification task.

In the present context, there is an analogue to the distributional hypothesis in that co-occurring *k-mers* are likely associated with the same or similar proteins, potentially sharing structure and function, and as we have seen, physico-chemical properties. We here build upon the earlier notion of context driven embeddings based on these *k-mers*, incorporating the available sequence labels to obtain sequence representations through a novel optimisation framework which we call `SuperVec`. As we shall discuss later, the `SuperVec` method performs better for the sequence retrieval task when compared with embeddings obtained using the earlier approaches, while retaining the computational efficiency critical when dealing with a large biological sequence database.

### 2.1 Word2Vec Architecture

We now briefly discuss the `Word2Vec` architecture, which forms a core building block for `SuperVec`. As discussed above, the idea behind `Word2Vec` is that pairs of words which share a semantic relationship should be proximally located in the embedding space. `Word2Vec` achieves these embeddings through the fully-connected shallow neural network (NN) shown in Fig 1. The input and the output layer of this network have *V* nodes, where *V* is the vocabulary size. Each node at the input/output layer is mapped to a vocabulary word such that the *i^th^* node corresponds to the *i^th^* word in the vocabulary. The weight vector **v**_*i*_ ∈ ℝ^*n*^ denotes connections from the *i^th^* input node to the hidden layer nodes, where n is the number of nodes in the hidden layer. The concatenation of input node weight vectors forms the input-hidden layer weight matrix, **W** ∈ ℝ^*V×n*^. Similarly, the hidden-output layer weight matrix is denoted **W**′ ∈ ℝ^*n×V*^.

**Fig 1.**
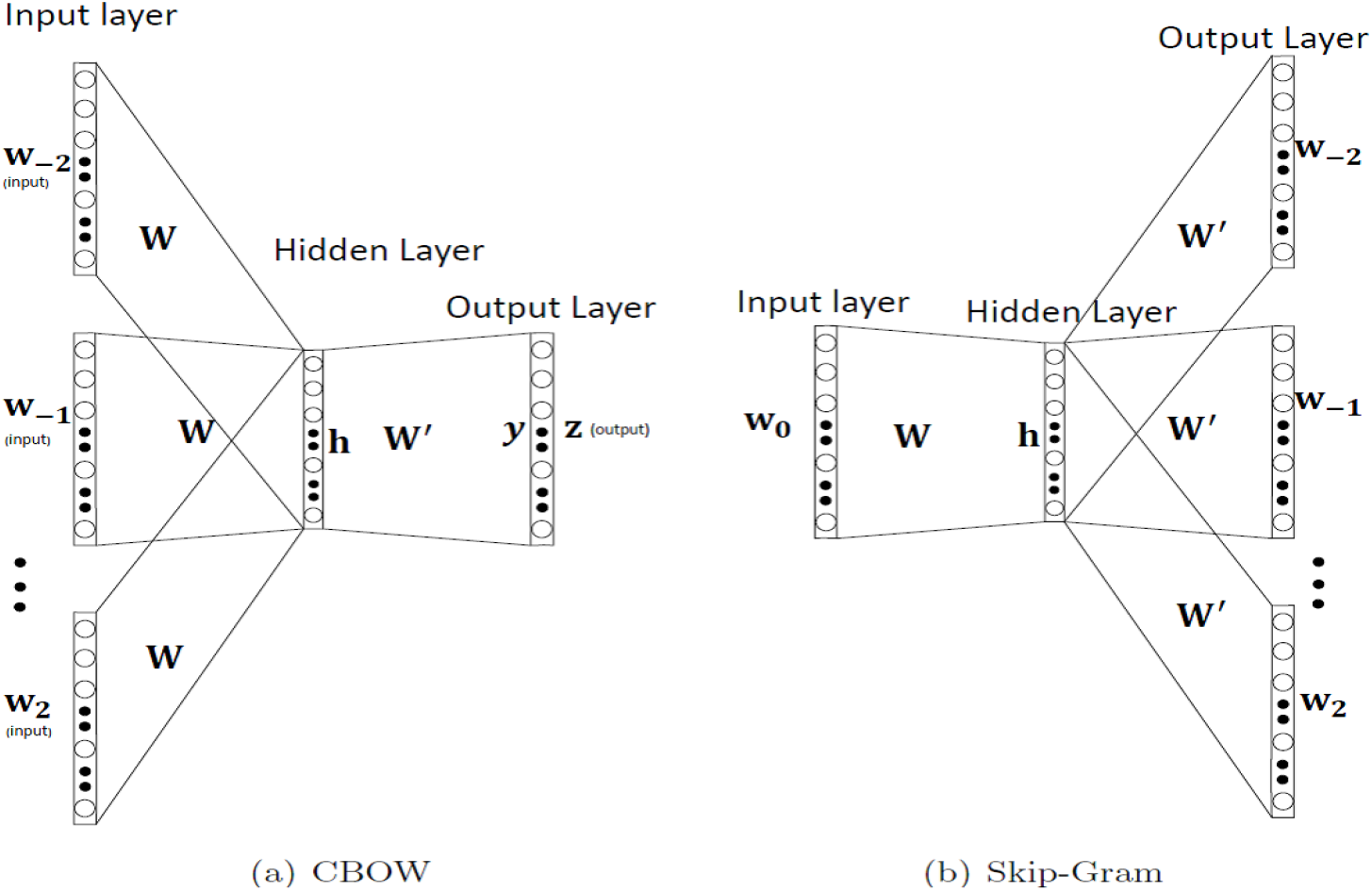
*Word2Vec* Architecture: The figure shows two variants of `Word2Vec` architecture - CBOW and Skip Gram [19]

Once training is completed, the weight vector **v**_*i*_ corresponding to the *i^th^* word at the input/output layer is also its corresponding embedding. These embeddings are learned such that words with semantically similar meanings are closer in the embedding space. This objective is achieved by a prediction based framework, where the loss function of the NN is a likelihood function for the words in the corpus. The likelihood function is proportional to the probability of the occurrence of a word given its context. The `Word2Vec` method has two variants, namely the CBOW (Continuous Bag of Words) and the Skip-gram. In the CBOW architecture, the context is given at the input to predict the word, whereas in the Skip-gram architecture, the word is used to predict the context.

Fig 1(a) shows a sample *w*_−2_, *w*_−1_, *w*_0_, *w*_1_, *w*_2_. Here, *w*_0_ denotes the word at the center and *w_i_* (*i* ≠ 0) denote the nearby context words. The subscript *i* denotes the position of words relative to the central word, *w*_0_. Inputs to the NN are here a one-hot-encoding of the context words, as shown in small bold letters in the figure.

A context is represented through a binary vector by keeping all elements zero except the indices corresponding to its constituent words. As shown in Fig 1(a), the one-hot-encoding of words in the context operates on the weight matrix **W** to give the hidden layer vector **h**, where

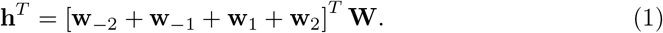

Here, **w**_−2_ + **w**_−1_ + **w**_1_ + **w**_2_ is the vector corresponding to the context. Similarly, the input to the output layer is given as **y**^*T*^ = **h**^*T*^**W**′. Finally the *Softmax* function is applied at the output layer to produce the output vector **z** = *softmax*(**y**). The output at node *j* is thus

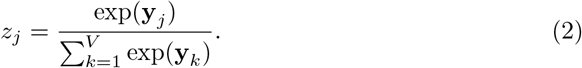

Here, *z_j_, y_j_* are the *j^th^* elements of the vectors **z** and **y** respectively. Vector **z** can be interpreted as the probability distribution of all words in the vocabulary given the context of *w*_0_ at the input. The embeddings for words in the corpus are obtained by maximizing the joint probability of a word, *w*, conditioned on its context, C, for all samples in the corpus. The conditional probabilities for each sample are assumed to be independent of each other and hence the joint probability for all samples can be written as, Π_*w*∈*Vocab*_ P*r*[*w*|*C*]. Maximizing Π_*w*∈*Vocab*_ P*r*[*w|C*] is the same as minimizing its negative log likelihood, so the overall loss function for `Word2Vec` may be written as

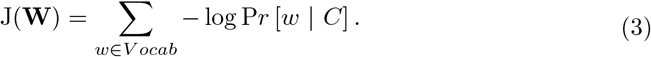

We note that updating the weight vectors implies adjusting those vectors corresponding to words in the embedding space. Maximizing the probability P*r*[*w*_0_|*C*], as given in Eq (2), is equivalent to maximizing the dot product of the weight vectors of *w_0_* and its context words, ultimately reducing the cosine distance between them. The process of maximizing the conditional probabilities (Eq (3)) therefore translates into reducing the cosine distance in the embedding space between the word and its surrounding context. The resulting optimisation problem can be solved using the standard gradient descent approach. However, to increase the computational efficiency of the algorithm, Mikolov et.al in [20] use a negative sampling method that essentially modifies the original objective function so that each training sample only updates a few weight vectors.

### 2.2 Doc2Vec Architecture

The `Doc2Vec` model essentially inherits the advantages offered by the `Word2Vec` model and generates a low dimensional representation of each document and its constituent words. These representations can then be used to retrieve documents from a collection. The main difference in the architecture of `Doc2Vec` and `Word2Vec` lies in the use of a document-tag along with the underlying words. The `Doc2Vec` architecture thus requires additional nodes to specify the document-tag as input. However, the output layer remains unchanged, and after training is completed, the `Doc2Vec` model yields both word embeddings and embeddings for the documents.

Having introduced the background to our work and the components which underpin our model, we now explain in detail the architecture and operation of `SuperVec`.

## 3 Proposed Approach: SuperVec

We now discuss our proposed supervised approach for training a sequence embedding framework, which once trained, can be utilised to generate embeddings for new unlabeled protein sequences. As outlined earlier, providing sequence labels while training may lead to a model that captures both a macro-level pattern from the class labels and the micro-level contextual information present in the protein sequences. We call our method `SuperVec`, reflecting the supervised nature of the approach. We discuss the model architecture and its operation in detail in following sections. We then introduce a new approach which utilizes the fact that `SuperVec` generates diverse and multiple embeddings for a sequence when trained with a diverse set of classes. These sets are obtained by partitioning the classes randomly and hence generating a tree-like structure (shown in Fig 3). The diversity of sequence embeddings gives us diverse query-subject distances which are processed jointly to give a better estimate of the similarity between the query-subject pair. We call this approach `Herierchical` `SuperVec` or `H-SuperVec`. We show in the result section that the use of label information during the training together with an increased diversity of representation models considerably increases the retrieval performance while continuing to offer good computational performance.

### 3.1 Notation

We denote the corpus of *N* protein sequences as 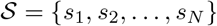 and the vocabulary with *M* unique *k-mers* as 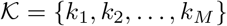. Each sequence *s_i_* = [*k*_*i*_1__, *k*_i_2__,…, *k_i_n_i___*] is an ordered list of *k-mers*. To avoid notation clutter we use *s_i_* to also denote its tag. Each sequence in 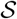 belongs to one of the *L* classes *l*_1_, *l*_2_,…,*l_L_* and *s_i_* ∈ *l_k_* means *l_k_* is the label for sequence *s_i_*. Finally, the embeddings of the *k-mer k_i_* and the sequence *s_i_* are denoted k_*i*_ and s_*i*_ respectively.

### 3.2 Training SuperVec

#### 3.2.1 Model Description

The `SuperVec` architecture consists of two `Word2Vec` units combined so that it can jointly incorporate the class label and contextual information contained in the sequences. As shown in Fig 2, NN1 is a CBOW configuration of `Word2Vec` that generates an embedding for sequences using the contextual information, while NN2 follows a skip-gram configuration of `Word2Vec` to help incorporate the class level information. NN1 requires a word-context pair for training while NN2 uses a sequence and a set of similarly labeled sequences. Note that NN1 forces the embeddings of the words that co-occur together along with the sequence embedding, while NN2 further constrains sequences having the same labels to be close to each other. Achieving embeddings informed *both* by context and by class information requires the use of both networks. Here we couple NN1 and NN2 by sharing the sequence embedding between them.

**Fig 2.**
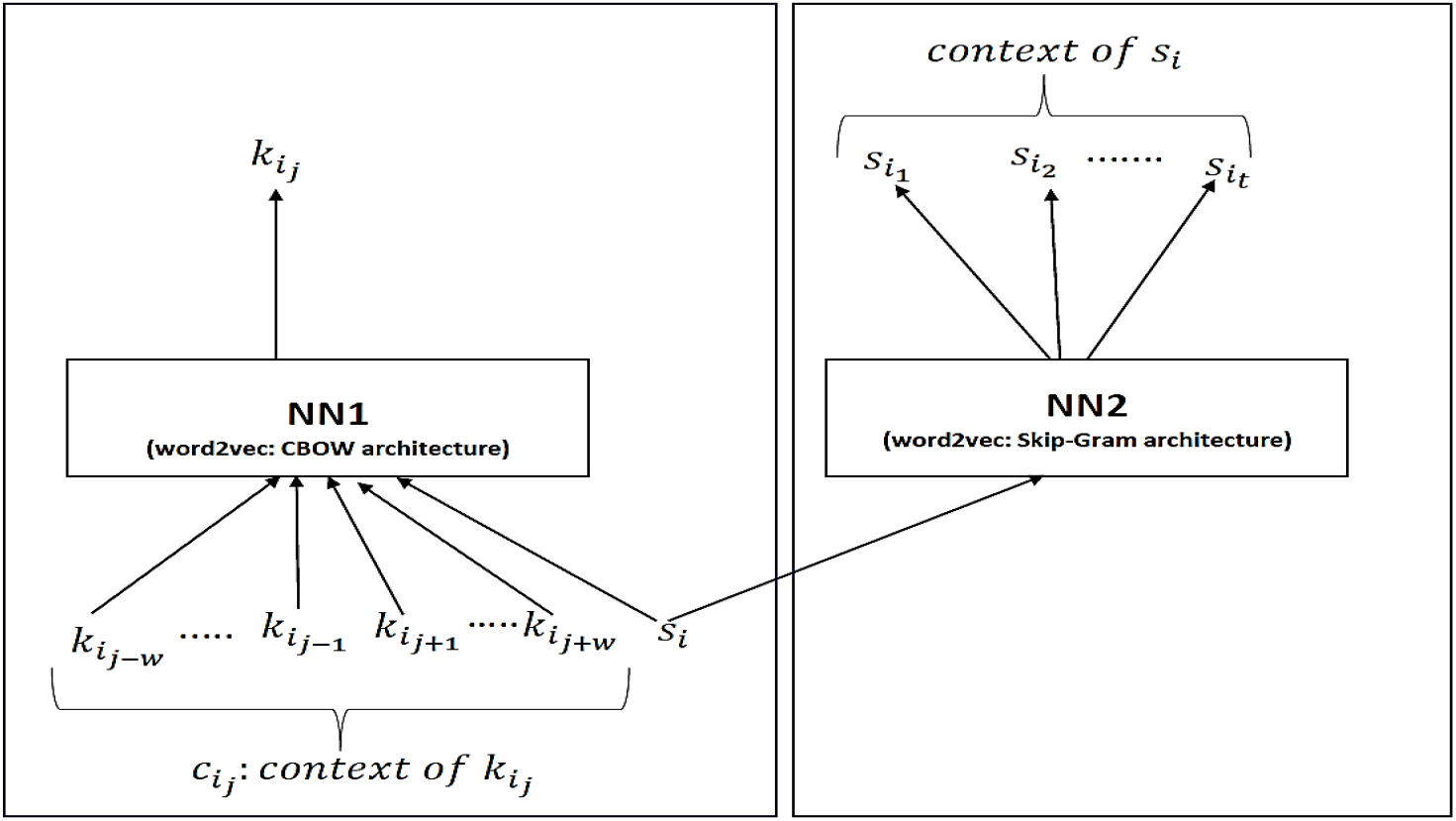
SuperVec Model: NN1 and NN2 architectures are shallow neural networks with respectively the CBOW and the Skip-gram variant of the `Word2Vec` model. The context of *k_i_j__*. comprises its nearby words and is denoted 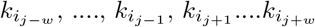. The context of the *s_i_* are sequences which have the same label as *s_i_*.

#### 3.2.2 Optimization problem formulation

To train the `SuperVec` model, i.e., to learn its parameters, two prediction tasks are performed, one by each of its sub-networks. Both networks essentially perform word(context)/context(word) prediction, although the meaning of context and word differs between them, as seen in Table 1 below. Here we see the sample *kmer k_i_j__*, its context *C_i_j__* and the corresponding sequence tag *s_i_*, and the relationships between them.

**Table 1.** context-word pairs for NN1 and NN2

| Network | Architecture | context | word |
| --- | --- | --- | --- |
| NN1 | CBOW | $s_i$ and $C_{i_j}$ | $k_{i_j}$ |
| NN2 | Skip-Gram | $s_{i_1}, s_{i_2} \dots s_{i_t} \in l_k$; given $s_i \in l_k$ | $s_i$ |

Since the sequence embedding task is shared by the sub-networks, the parameters of these networks influence each other. Mathematically, the coupling of these sub-networks means solving a joint optimization problem with the overall loss function a linear combination of those for NN1 and NN2

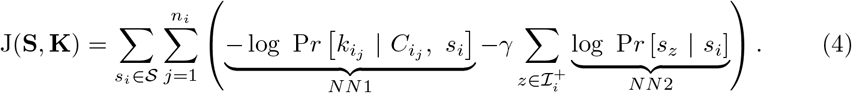

Here *γ* controls the balance between class information and contextual information. The matrices **K** = [**k**_1_, **k**_2_,…,**k**_*M*_]^*T*^ ∈ ℝ^*M × n*^ and **S** = [**s**_1_, **s**_2_,…,**s**_*N*_]^*T*^ ∈ ℝ^*N×n*^ denote the embeddings for sequences and *k-mers* respectively. We define 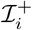 to be the set of indices of sequences which have the same label as *s_i_*. The conditional probability for NN1 in Eq (4) can be computed by a softmax function, but considering the large number of *k-mers* and the computational burden involved, we approximate it using hierarchical softmax (HS) [21]. To compute the second part of Eq (4) we use *negative sampling* [20], where for any given sequence we try to maximize the probability of some selected positive samples as opposed to others. For example in a context/word pair, all the words in the context can be treated as positive samples and the remaining words in the vocabulary as negative samples. In our case, for any input sequence *s_i_*, sequences whose index lies in 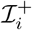 are seen as positive samples, whereas the remaining sequences are seen as negative samples. We denote the set of indices of the negative samples as 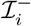. The second part of Eq (4) can then be approximated as

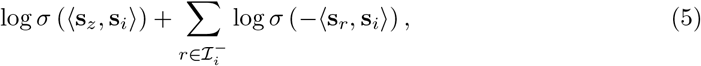

where 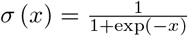 is the sigmoid function. Maximizing the first part of the Eq (5) maximizes 〈s_*z*_, s_*i*_〉 and therefore reduces their cosine distance in the embedding space. Similarly, maximizing *σ* (−〈s_*r*_, s_*i*_〉) translates into maximizing the cosine distance between s_*i*_ and s_*r*_. As defined above, the *s_z_* have the same label as *s_i_* whereas the *s_r_* have a different label. Maximizing Eq (5) therefore forces the sequences from the same class to be mapped closer in the embedding space, while other sequence pairs are pushed apart. Employing negative sampling thus yields embeddings with low intra-class and high inter-class separation, a mapping well suited to the retrieval task. Replacing the second part of Eq (4) with its *negative sampling* expansion, the final loss function for `SuperVec` is given as

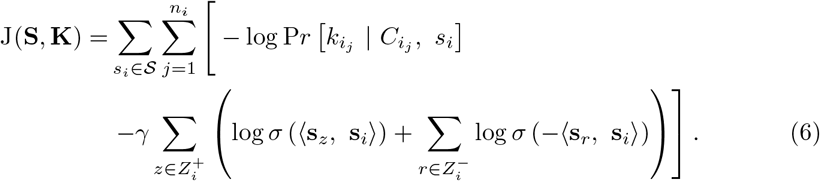

Computing the second part of Eq (6) requires a very large number of operations—the *number of positive samples × number of negative samples*—for each case. To mitigate this computational burden, we use only a few randomly selected positive and negative samples for any given sequence. In Eq (6), 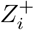 and 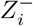 are sets which constitute the indices randomly chosen from the positive indices, 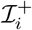, and from the negative indices, 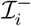, respectively. Note that we sample positive as well as negative samples to further speed up the training process.

#### 3.2.3 Parameter learning

Before discussing the parameter learning process for `SuperVec`, it is important to note that as for the `Word2Vec` model, the parameters of `SuperVec` constitute the embeddings of the *k-mers* and the sequences. At the end of the training process, we not only get a trained network but we also obtain the embeddings for *k-mers* and the sequences used in the training process. The process of learning the parameters involves a training process similar to that used for the `Word2Vec` framework. The parameters are initialized randomly and then modified for each sample to reduce the value of the loss function. The sample here consists of the *k-mer*, its context, and the corresponding sequence tag. As explained before, `SuperVec` employs two prediction tasks for each selected sample. In these prediction tasks, for any context/word at the input, the parameters are modified to maximize the probability of word/context at the output by updating the values of parameters using gradient descent. After sufficient iterations, we obtain a trained network which retains the relevant information – in this case, the contextual and class information. The update equations and the derivation of the gradient for s_*i*_, k_*i_j_*_ and other parameters of `SuperVec` over **J**(**S, K**) are provided in the appendix ??. Once the model is trained it can be employed for learning the representation of any new sequence. Note that new sequences do not require any label information: as discussed below, we use only NN1 for learning this representation.

#### 3.2.4 Inference

Computing the representation of a new sequence by passing it through the trained model constitutes the inference step, and a model which generates meaningful representations for new sequences may prove useful for many downstream bioinformatics tasks. The efficacy of the inference step is evaluated with respect to the task for which the learned representations are employed. In this paper, we consider retrieval as the downstream application. Here it is expected that the information learned by the trained model is exploited in the representation of the new sequence.

For the retrieval task, `SuperVec` is trained over labeled sequences. Once we have the trained `SuperVec` model, we use it to generate embeddings for the database sequences. When a new sequence is given as a query, the first step is to generate its representation within the embedding space, and to compute the relevant distances. Since the inference step does not utilize the sequence labels, only one of the sub-networks of `SuperVec`, i.e. NN1 (refer to Fig 2), is used. Although the standalone NN1 only uses contextual information, we expect that since NN1 and NN2 are coupled and jointly trained, NN1 will also capture the class information to some extent, and that this will eventually influence the representation learned in the inference step. While the representation for a sequence is computed during the inference step, all the parameters of NN1 remain unchanged except for the sequence vector, which is initialized with zeros. This vector is updated iteratively following a gradient descent approach similar to the training stage.

The comparison of `Seq2Vec` and `SuperVec` results for retrieval tasks confirms our hypothesis that incorporating label information in addition to the contextual information in the training process helps in generating useful sequence embeddings for retrieval purposes. Although `SuperVec` performs better than other representation learning methods (`Seq2Vec` [7], `BioVec` [6]) for a number of retrieval tasks, we observe that its retrieval performance deteriorates with an increasing number of classes. With the increase in the number of classes, the constraints enforcing interclass and intraclass separation necessarily increase in number. Satisfying this set of constraints may prove difficult, reducing the efficacy of the training process and ultimately leading to a deterioration in retrieval performance. The other important observation to note here is that the interclass and intraclass distances of a set of sequences change when we generate sequence embeddings using `SuperVec` models trained on a diverse set of classes. This observation implies that we get a diverse embedding of a sequence when generated through multiple models. Keeping these observations in mind, we propose a hierarchical approach which improves the retrieval results by computing a better estimate of the query-subject similarity. We call this proposed method `H-SuperVec`. In this approach we work with a binary-tree obtained by partitioning a set of classes at each parent node (refer Fig 3). Once the tree is created, a `SuperVec` model is trained for each of the nodes. Following this approach gives us multiple trained models, which can be used to generate many observations of the same quantity (here the query-subject distance). These observations are processed jointly to get a better estimate of query-subject similarity. We describe the `H-SuperVec` model in detail in the following section.

**Fig 3.**
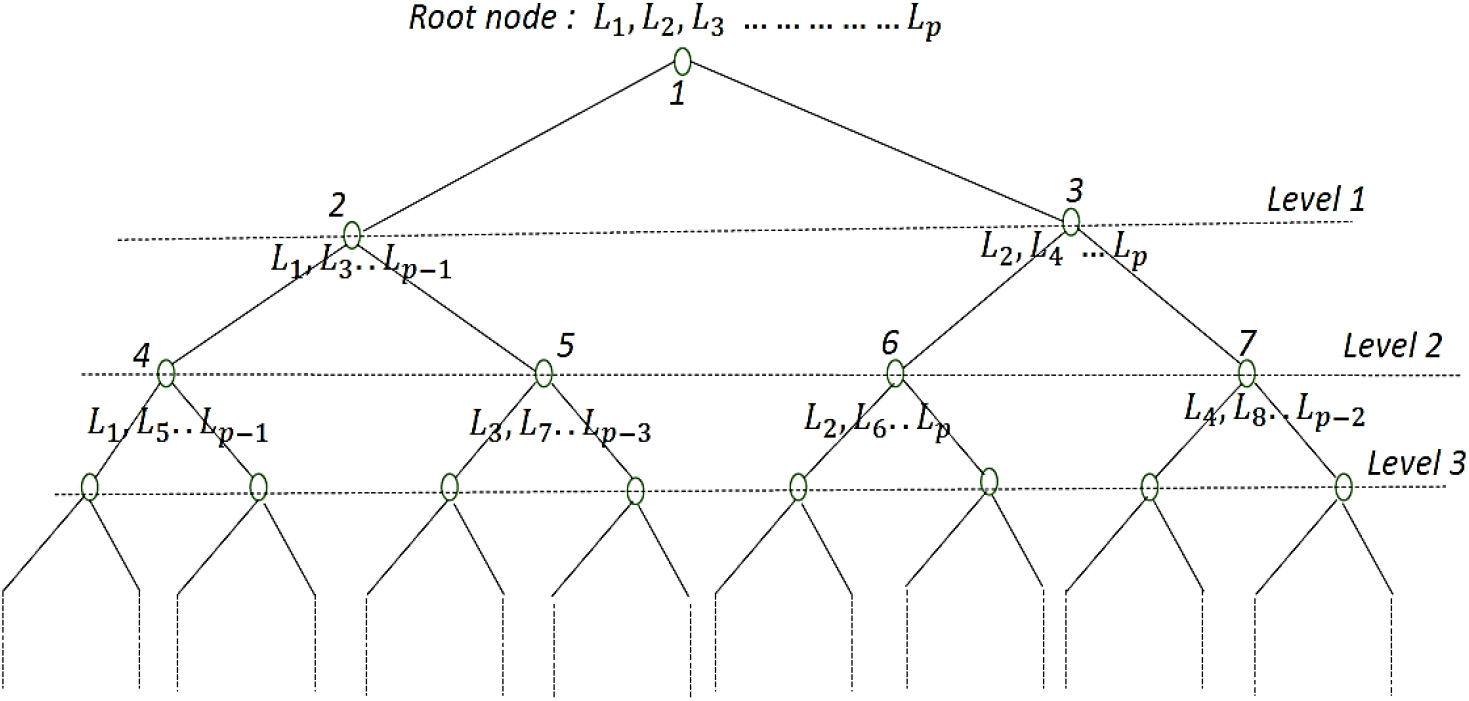
An example Binary tree for H-SuperVec: A hierarchical structure obtained by partitioning the class labels of each parent node into equal size subsets. The root node is assigned *p* class labels (*L*_1_, *L*_2_….*L_p_*) and their corresponding sequences. In this example we assume that p is an even number; the right child of the parent node is assigned the even and the left child is assigned the odd indexed labels selected from those assigned to the parent node.

### 3.3 Hierarchical SuperVec

`H-SuperVec` exploits the fact that we can generate multiple, diverse embeddings for a given sequence using multiple `SuperVec` models trained on sequence data belonging to a diverse set of classes. Note that for each query-subject (database sequence) pair, each `SuperVec` model results in a distance computation corresponding to that model. This distance is used as a proxy to measure the similarity between two sequences. Each `SuperVec` model introduces some noise in the embeddings it generates and hence in the distance computed for any query-subject pair. Processing the query-subject distances obtained from different `SuperVec` models together reduces the overall noise and give us a better estimate of query-subject similarity. We utilize this fact in `H-SuperVec` and apply it to the same retrieval task. Applying `H-SuperVec` for retrieval tasks involves the following steps:

- Form a Hierarchical Structure: First, we assign all of the class labels and their corresponding sequences to the root node. The root node is then split into two child nodes by randomly partitioning its associated class labels into equal halves. These child nodes are further partitioned, following the same process for each node until we are left with leaf nodes, each associated with a single class label. An example of such an hierarchical tree is shown in Fig 3.
- Train a SuperVec model for each node of the above tree: these models can subsequently be used to generate embeddings for a new sequence.
- Assign weights to each node: As we traverse down the tree, `SuperVec` is successively trained with fewer classes, leading to an increase in noise in the computed query-subject distance. To get a better estimate of query-subject similarity, we apply a simple linear model (weighted sum) over the distances computed at each node. For query (*q*), the similarity is estimated as the weighted sum:

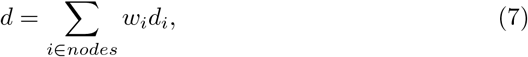

where *d_i_* is the query-subject distance computed at the *i^th^* node and *w_i_* is the weight assigned to node *i*. Since noise increases as we traverse down the tree, the largest weight is assigned to the root node, with node weights decreasing as we traverse toward the leaves. The weight magnitudes are constrained by three conditions: (i) the weights are positive, *w_i_* > 0, ∀*i*; (ii) nodes at the same level of the tree are assigned equal weights; and (iii) the weights sum to one, i.e., Σ_*i*∈*nodes*_ *w_i_* = 1.
- Retrieve sequences: Once we build a hierarchical tree and train the `SuperVec` model for each node, the retrieval task is performed as follows. First, we learn embeddings for database sequences using the `SuperVec` model at each node. For a new query sequence, multiple embeddings are generated corresponding to the nodes of the tree using the inference step. Pairwise query-subject distances are then calculated for each node using Eq (8). These distances are finally combined as a linear sum to give an estimate of similarity between the query-subject pair. The results are returned in descending order of their similarity with the query sequence.

Since the weight assigned to each node is reduced as we traverse down the tree, the contribution of the nodes in the computation of similarity between query-subject pair also decreases. We empirically determined that working with a one or two level tree produces consistently better retrieval results. For our experiments, we chose a tree with only one level, i.e., the tree having root and its child nodes. We demonstrate the mechanism followed by the `H-SuperVec` method to estimate the distance of query-subject pair in Fig 4. `SuperVec1`, `SuperVec2` and `SuperVec3` are the models for root and first level nodes respectively. Each of these models is utilized to obtain the query-subject distance; the distance computed at *i^th^* node and hence through *i^th^* model is given as,

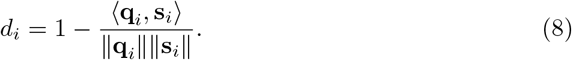

where **q**_*i*_ and s_*i*_ are the embeddings of query ‘q’ and a database sequence ‘s’ computed at *i^th^* node. The weights *w_i_* were chosen empirically in order to optimise retrieval performance subject to the constraints discussed above, and in this initial study we have limited experiments to shallow trees, where the depth does not exceed 3. Retrieval performance is generally stronger if the weight mass is concentrated toward the root node, with the best results obtained with the set {0.75, 0.125, 0.125}, corresponding to root and child nodes respectively. Employing `H-SuperVec` for the retrieval task gives an improvement over all representation learning method considered in this paper including `SuperVec`; the results are shown in section 5.

**Fig 4.**
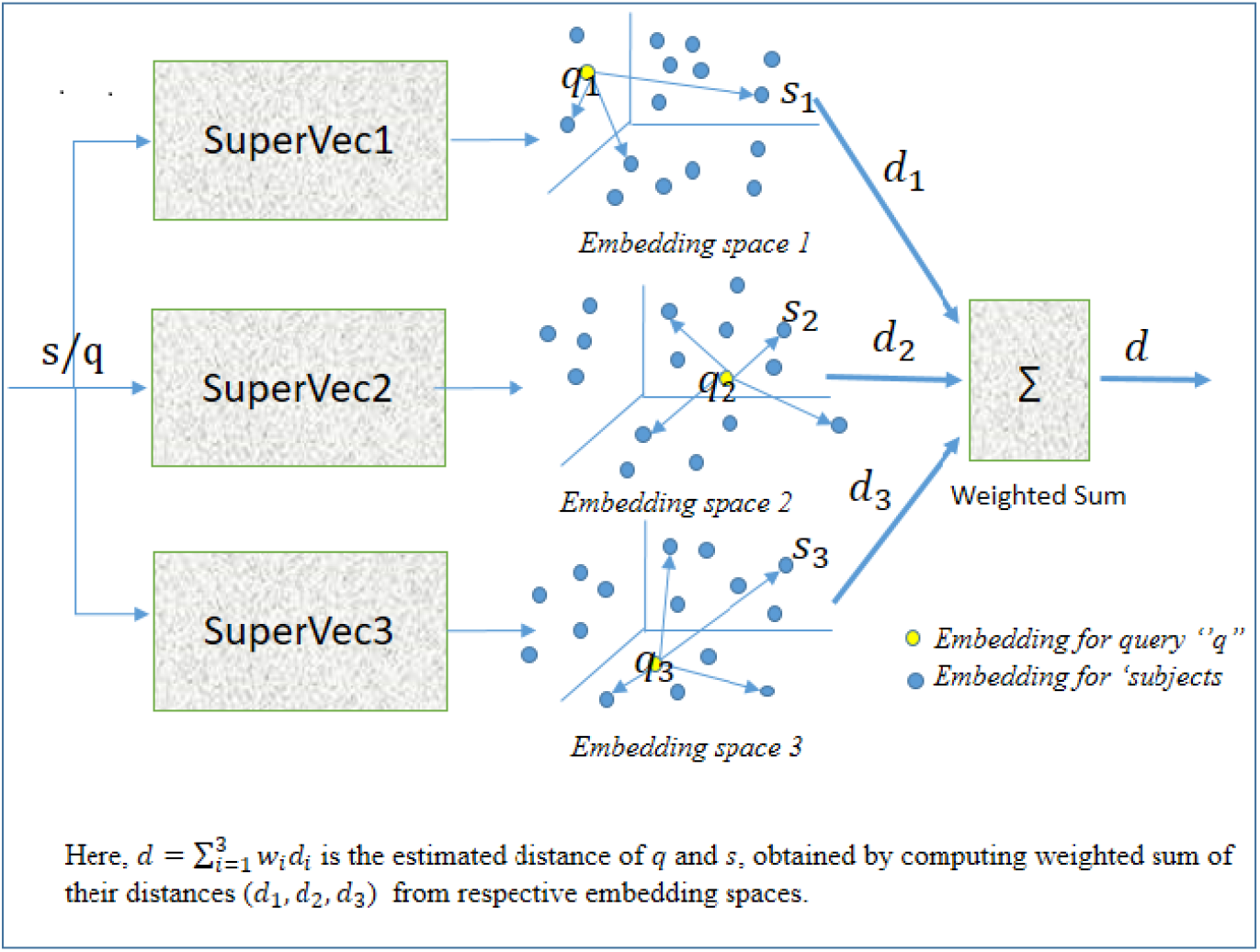
`SuperVec`1/2/3 are the `SuperVec` model trained for node 1, 2 and 3 respectively. **q**_*i*_ and **s**_*i*_ are the embedding of a query (*q*) and subject (*s*) and *d_i_* is the distance of query-subject pair computed at *i^th^* node, *d* is the similarity score for *q* and *s*.

## 4 Experimental Design and Evaluation

### 4.1 Task and Data

We now discuss the experimental setup to demonstrate the quality of embeddings generated using the proposed method. Asgari et. al [6] demonstrated the utility of the embeddings generated through their method `BioVec` on the protein family classification task. In this work, we focus on the related problem of homologous sequence retrieval, where the task is to return a set of sequences from a database that are homologous to a given query. For embedding based methods, we address the retrieval problem by employing nearest neighbour search, where for a given query the sequences from a database are retrieved based on their cosine-distance to the query sequence embedding.

We use the protein sequence data provided by Asgari et al. [6] for evaluation of our approach on the sequence retrieval task. The dataset consist of 324018 protein sequences, each uniquely annotated with one of the 7027 family labels. In this dataset, there are families who have the same functional description given in the PFAM database. For example, Chitin_synth_1 (PF01644) and Chitin_synth_2 (PF03142) are two different entries in PFAM, but both represent the chitin synthase enzyme. We merged such families into a single representative family; this led to a reduction in the number of families to 6967. The families/classes present in this dataset differ considerably in their size; the largest family contains 3024 sequences, whereas many families are based on a single sequence. The distribution of class sizes and sequence lengths is shown in Fig 5.

To create reasonably sized training and test splits for experiments we chose the largest 200 families, ensuring a minimum family size of least 400 members. These families contain a total of 15, 0324 sequences.

**Fig 5.** Distribution of class sizes and lengths of sequences present in the complete dataset.

In the experiments below, we first demonstrate the utility of our method on a small (two-class) data-set. Subsequently, we perform retrieval experiments on different database sizes ranging from 21531 (25 classes) to 90121 sequences (200 classes), to analyze the reliability of our method.

### 4.2 Experimental Design

The experimental setup used to conduct most of the retrieval experiments in this paper is discussed below.

- **Setup 1:** This setup was designed to demonstrate the advantages of the supervised method `SuperVec` and its extension `HSuperVec` on the sequence retrieval task. Here, the database was constructed by choosing at random 60% of the sequences from each class (each family corresponds to one class). The remaining 40% of these sequences form a query set that is used to evaluate retrieval performance on the database. Retrieval based on a given query proceeds as follows. First, for each database sequence, an *n*-dimensional embedding is generated, yielding a database embedding space. For a new query, we first generate its embedding; then we employ nearest-neighbor search in the database embedding space, returning sequences in descending order of cosine similarity. To avoid the computational overburden of computing the distance of all database sequences to the query and sorting them, we use an approximate neighborhood search [22] technique and fix the neighborhood size to be 10, 000. Note that the process of generating embeddings for the sequences differs for each of the considered RL methods. For `BioVec`, sequence embeddings are generated by adding the corresponding *k-mer* embeddings; we use the *k-mer* embeddings provided by [6]. To generate the sequence embedding using the `Seq2Vec` or `SuperVec` approach, we first train them using the database sequences; note that unlike `Seq2Vec, SuperVec` also uses the database sequence label for training. Once these models are trained, the sequence vectors are generated by following the inference step.

In this study, all experiments were performed using a commodity Linux workstation equipped with an Intel Core i7-4790K, 3.6GHz 8 core, 16 thread processor. The values of hyper-parameters–the *k*-mer size and the representation length of sequences for `SuperVec` and `Seq2Vec`–are kept same as given in [6], whereas the context size and supervision parameter *γ* are chosen based on the best retrieval results obtained on a database of largest 50 classes under setup-1. The values selected for the `SuperVec` and `Seq2Vec` hyper-parameters are shown in Table 2.

**Table 2.**
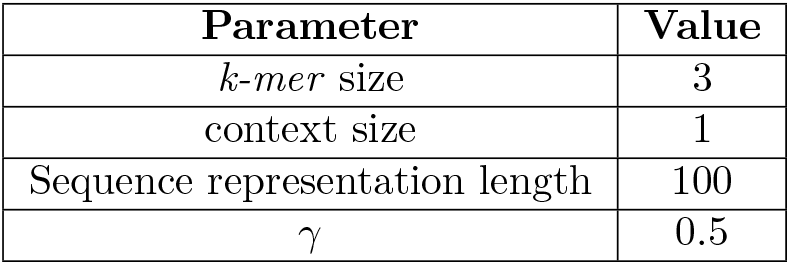
Hyper-parameters for `SuperVec/Seq2Vec`

### 4.3 Evaluation

The performance of `SuperVec` and `H-SuperVec` on the homologous sequence retrieval task was evaluated through comparison against the other representation learning (RL) approaches–`BioVec` and `Seq2Vec`–along with BLAST, the standard approach for this problem.

The evaluation metrics for the retrieval task were chosen as follows:

1. **Interpolated precision-recall values:** The interpolated precision value *p*′(*r*) [23] at a recall level *r*, is defined as highest precision value found for any recall level *r*′ > *r*; 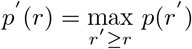. Here, *p*(*r*) is the precision value at recall *r*, which is defined as the ratio of the number of relevant sequences to be retrieved at recall *r*, and the number of sequences that need to be retrieved to get that many relevant sequences for a given query.
2. **Querying Time:** The querying time is defined as the time required to retrieve the significant matches from the database for a given query. As discussed before, the retrieval process for RL methods requires that we first train the model, which is then used for generating the representations for the database and subsequent query sequences followed by the nearest-neighbor search. This training process is computationally intensive, but is required only once for each model. The training time for `SuperVec` for ~ 90*k* training sequences is ~ 28 hours, and there remains scope to improve this performance through parallelization and optimization. We compute the querying time as the sum of the time taken to generate the sequence embedding and the time required to produce the list of nearest neighbor(s) for a given query. For BLAST, we report the querying time as the time required for it to return the list of database sequences for a given query, i.e. we do not consider the creation of the BLAST database.

## 5 Results and Discussion

In this section, we present the results obtained on the retrieval task for `SuperVec, H-SuperVec` and the other methods considered.

### 5.1 Supervised Vs Unsupervised Embeddings

Biological sequence embeddings generated through Representation Learning methods can be broadly categorized as supervised or unsupervised based on the training process adopted. Here, `SuperVec` will generate supervised embeddings, while approaches such as `BioVec` and `Seq2Vec` will yield unsupervised embeddings. The information captured in these embeddings plays a vital role in ensuring a suitable basis for the task undertaken. In the case of retrieval, high precision requires that the representation: (i) generate a database embedding space with low intraclass and high interclass separation, and (ii) map each new query close to its class in the embedding space.

To demonstrate the effect of the supervision adopted in `SuperVec` on database and query embeddings, and its consequences for retrieval performance, we initially consider a retrieval task over a small database – consisting of the two largest classes from the dataset. We then visualize the two-dimensional t-SNE plots [24] of the database and query sequence embeddings generated through `SuperVec, BioVec` and `Seq2Vec` and subsequently compare their retrieval performance.

From the t-SNE plots shown in Fig 6, we observe that:

- `BioVec` generated database embeddings are well-separated by class, but form small groups within each class. `Seq2Vec` provides (relatively) better intraclass organisation than `BioVec`, but the two classes are merged to a great extent. `SuperVec` database embeddings show a better intraclass and interclass separation than the other methods, albeit with some overlap at the boundary.
- For a new query, the presence of the relevant subjects (here the database sequences from the same class) decreases as we increase the neighborhood size. Thus, we expect to see a decrease in the precision values for increasing recall levels. Analyzing the plots we can infer that such an effect will have a stronger impact on the retrieval performance of `Seq2Vec` and `BioVec` when compared to `SuperVec`, especially for late recall levels. To validate this observation, we compare the retrieval performance of the RL methods considered, and find results consistent with our observations from the t-SNE plots. From the retrieval results in Table 3, we can infer that supervision helps in the retrieval task.
- To ensure that these results hold more broadly, we conducted a similar two-class experiment for 100 randomly selected pairs and found the outcomes consistent with the largest pairing discussed above. The results are shown in Table 4. We also conducted same experiments on the largest eight and sixteen classes and observe that `SuperVec` consistently outperforms other RL methods. The results are shown in Table 5 and Table 6 respectively.

**Fig 6.**
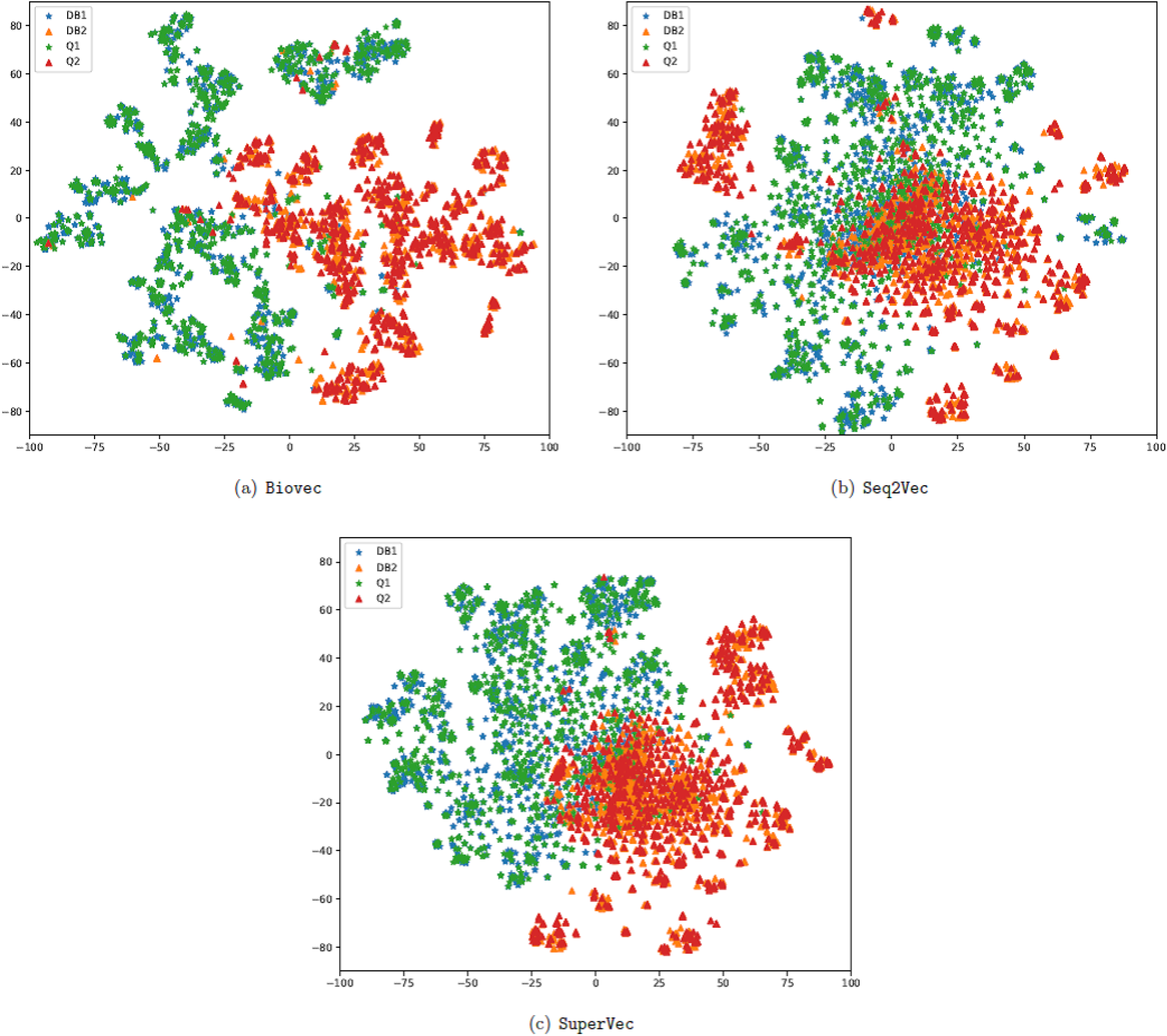
t-SNE plots: The mapping of database and query embeddings generated through `BioVec, Seq2Vec` and `SuperVec` approaches for largest two classes from the dataset; DB1, DB2 denotes the database sequences and Q1, Q2 denotes the query sequences from class 1 and class 2 respectively. There is a better segregation of the DB1 and DB2 for `SuperVec` as compared to `Seq2Vec` and `BioVec`. Also, the presence of number of relevant database sequences for a given query decreases at a lower rate for `SuperVec` followed by `BioVec` and `Seq2Vec` as we increase the neighborhood size.

**Table 3.**
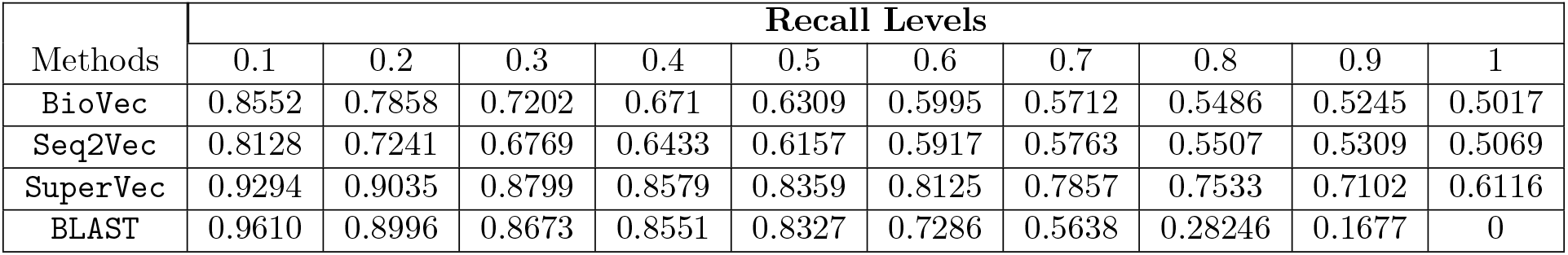
Largest two class experiment results: Average interpolated precision values at ten recall levels computed for 2242 sequences queried on the database of largest two classes, 3360 sequences. The average precision value at a particular recall level is calculated by first computing the class wise average followed by averaging the precision value obtained for both the classes.

| Methods | Recall Levels |  |  |  |  |  |  |  |  |  |
| --- | --- | --- | --- | --- | --- | --- | --- | --- | --- | --- |
|  | 0.1 | 0.2 | 0.3 | 0.4 | 0.5 | 0.6 | 0.7 | 0.8 | 0.9 | 1 |
| **BioVec** | 0.8552 | 0.7858 | 0.7202 | 0.671 | 0.6309 | 0.5995 | 0.5712 | 0.5486 | 0.5245 | 0.5017 |
| **Seq2Vec** | 0.8128 | 0.7241 | 0.6769 | 0.6433 | 0.6157 | 0.5917 | 0.5763 | 0.5507 | 0.5309 | 0.5069 |
| **SuperVec** | 0.9294 | 0.9035 | 0.8799 | 0.8579 | 0.8359 | 0.8125 | 0.7857 | 0.7533 | 0.7102 | 0.6116 |
| **BLAST** | 0.9610 | 0.8996 | 0.8673 | 0.8551 | 0.8327 | 0.7286 | 0.5638 | 0.28246 | 0.1677 | 0 |

**Table 4.** Retrieval results for 100 random pairs: Average interpolated precision values at ten recall levels computed for 100 random pair of classes. All of these pairs differ in the number of database and query sequences. The precision value shown at particular recall level below is averaged over the chosen 100 pairs.

| Methods | Recall Levels |  |  |  |  |
| --- | --- | --- | --- | --- | --- |
|  | 0.1 | 0.2 | 0.3 | 0.4 | 0.5 |
| BioVec | $0.9079 \pm 0.087$ | $0.8817 \pm 0.101$ | $0.8573 \pm 0.112$ | $0.8329 \pm 0.124$ | $0.8066 \pm 0.134$ |
| Seq2Vec | $0.9494 \pm 0.034$ | $0.9258 \pm 0.05$ | $0.904 \pm 0.064$ | $0.8828 \pm 0.076$ | $0.8616 \pm 0.085$ |
| SuperVec | $0.9899 \pm 0.009$ | $0.9882 \pm 0.011$ | $0.9868 \pm 0.013$ | $0.9853 \pm 0.014$ | $0.9836 \pm 0.016$ |
| BLAST | $0.9705 \pm 0.046$ | $0.9562 \pm 0.082$ | $0.9416 \pm 0.105$ | $0.9322 \pm 0.115$ | $0.9208 \pm 0.133$ |
|  | 0.6 | 0.7 | 0.8 | 0.9 | 1 |
| BioVec | $0.7764 \pm 0.142$ | $0.7458 \pm 0.147$ | $0.7109 \pm 0.149$ | $0.6617 \pm 0.143$ | $0.5520 \pm 0.082$ |
| Seq2Vec | $0.8380 \pm 0.094$ | $0.8110 \pm 0.102$ | $0.7778 \pm 0.108$ | $0.7320 \pm 0.111$ | $0.6247 \pm 0.095$ |
| SuperVec | $0.9815 \pm 0.018$ | $0.9788 \pm 0.021$ | $0.9749 \pm 0.025$ | $0.9675 \pm 0.032$ | $0.9279 \pm 0.06$ |
| BLAST | $0.9084 \pm 0.153$ | $0.8918 \pm 0.180$ | $0.8547 \pm 0.214$ | $0.7896 \pm 0.259$ | $0.0485 \pm 0.136$ |

**Table 5.** Largest eight class experiment: Average interpolated precision values at ten recall levels computed for 6553 sequences queried on the database of largest eight classes, 9793 sequences. The average precision value at a particular recall level is calculated by first computing the class wise average followed by averaging the precision value obtained for all the classes.

| Methods | Recall Levels |  |  |  |  |  |  |  |  |  |
| --- | --- | --- | --- | --- | --- | --- | --- | --- | --- | --- |
|  | 0.1 | 0.2 | 0.3 | 0.4 | 0.5 | 0.6 | 0.7 | 0.8 | 0.9 | 1 |
| BioVec | 0.7168 | 0.623 | 0.56 | 0.5023 | 0.4265 | 0.3592 | 0.3119 | 0.2667 | 0.2218 | 0.1134 |
| Seq2Vec | 0.6671 | 0.5604 | 0.4886 | 0.4374 | 0.3863 | 0.3448 | 0.3079 | 0.2719 | 0.2335 | 0.1508 |
| SuperVec | 0.7985 | 0.7437 | 0.7016 | 0.6667 | 0.6342 | 0.5998 | 0.5571 | 0.4783 | 0.379 | 0.1799 |
| BLAST | 0.9631 | 0.9348 | 0.9110 | 0.8918 | 0.8307 | 0.7382 | 0.66 | 0.4716 | 0.337 | 0 |

**Table 6.** Largest sixteen class experiment: Average interpolated precision values at ten recall levels computed for 10537 sequences queried on the database of largest two classes, 15791 sequences. The average precision value at a particular recall level is calculated by first computing the class wise average followed by averaging the precision value obtained for all the classes.

| Methods | Recall Levels |  |  |  |  |  |  |  |  |  |
| --- | --- | --- | --- | --- | --- | --- | --- | --- | --- | --- |
|  | 0.1 | 0.2 | 0.3 | 0.4 | 0.5 | 0.6 | 0.7 | 0.8 | 0.9 | 1 |
| BioVec | 0.7128 | 0.598 | 0.5304 | 0.4757 | 0.4154 | 0.3656 | 0.3229 | 0.258 | 0.178 | 0.0735 |
| Seq2Vec | 0.6684 | 0.5545 | 0.4857 | 0.4323 | 0.3837 | 0.3381 | 0.2901 | 0.22 | 0.1526 | 0.0808 |
| SuperVec | 0.8217 | 0.7782 | 0.744 | 0.7121 | 0.6771 | 0.6386 | 0.5935 | 0.5387 | 0.4641 | 0.1198 |
| BLAST | 0.9547 | 0.924 | 0.8985 | 0.8799 | 0.8341 | 0.7474 | 0.6815 | 0.5664 | 0.4662 | 0 |

As the RL approaches fall broadly under the umbrella of alignment-free methods, we also consider the performance of BLAST on these tasks a most widely used of the alignment-based approaches. The results show that `SuperVec` outperforms BLAST for the two class database problems, while the performance declines as the number of classes in the database increases. As we shall see later, we improve upon `SuperVec` through the use of `H-SuperVec` and `H-SuperVec+BLAST`. Comparing the querying time for each approach, we see that the RL methods provide a substantially faster alternative, offering a speedup in querying time of more than 40 × when compared to BLAST. For the experiments involving the largest eight and sixteen classes, the average querying time is ~ 20 – 30 ms for RL approaches and ~ 900 ms for BLAST.

These experiments were subsequently extended to much larger databases involving a large number of classes and sequences. We again followed experimental setup-1 and we limited our analysis to the top 200 classes, ensuring a minimum size of 400 samples. The results of these experiments are shown in Fig 7. The plots in Fig 7(a) and Fig 7(b) shows the interpolated precision-recall graph obtained for a small database–21531 sequences, 25 classes-and for a larger database–90121 sequences 200 classes-respectively. These results show that `SuperVec` maintains its superiority when compared to the unsupervised methods even for larger databases. Note that although the results for only two database sizes are provided in Fig 7, we performed the same retrieval task on the database of different sizes —35088 sequences 50 classes; 57768 sequences, 100 classes and 75576 sequences, 150 classes and find that the results remain consistent with the experiments shown.

**Fig 7.**
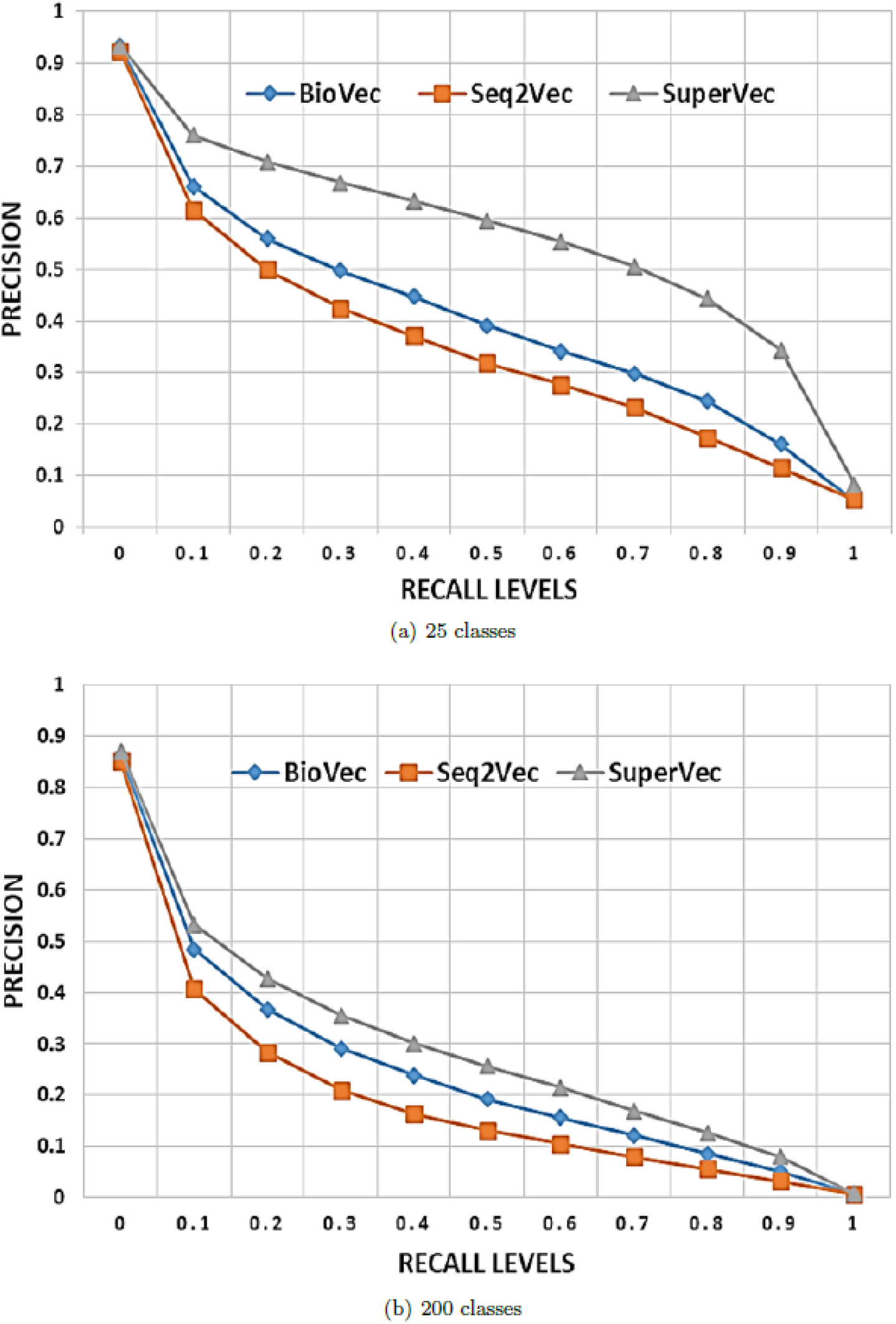
Supervised Vs unsupervised: Average interpolated precision values at 11 recall levels for largest 25 and 200 classes.

Our observations may be summarized as follows:

- `SuperVec` consistently outperforms the baseline biological sequence embedding approaches on the sequence retrieval task for a wide range of database sizes.
- RL methods are substantially faster than BLAST and provide a gain of ~35× in querying time, albeit with some reduction in precision.
- The presence of a large number of sequences and classes in the database leads to deterioration in the performance of all methods. Although the retrieval performance of `SuperVec` also deteriorates for larger databases it still consistently performs better than `Seq2Vec` and `BioVec`, most likely as a result of its ability to incorporate the class label information in the sequence embeddings.

#### 5.1.1 Robustness of SuperVec

To study the robustness of our approach, we conducted experiments on larger databases, this time following the different setup discussed below.

- **Setup 2:** In this setup the sequences are chosen randomly from each class in the ratio 60:20:20, generating the training, database and query set. Here the training sequences are used for training the models whereas the database and query sets are reserved for validation on the retrieval task. Once the models are trained, the process followed to generate sequence embeddings and retrieval of homologous sequences for a given query is the same as described in setup-1. Note that this change won’t have much effect on `BioVec` generated sequence embeddings as we use the *k-mer* embeddings directly from [6]. As mentioned before, setup-1 uses database sequences and their labels as part of the training process, leading to the possibility of overfitting with respect to the database sets. This setup is intended to show the robustness of our approach, demonstrating that our method gives retrieval results similar to those obtained in setup-1 even when a different set of sampled sequences are used to train the models.

The retrieval results obtained following setup 2 are consistent with setup-1, in which the supervised approach outperforms the baseline RL approaches. The retrieval result for the database of 100 class and 75576 sequences is shown in Fig 8. Since the database and training sequences are different in setup-2, the superior results obtained using `SuperVec` suggest that `SuperVec` transfers the information learned from the training sequence and their labels more efficiently to the database and query sequence embeddings.

**Fig 8.**
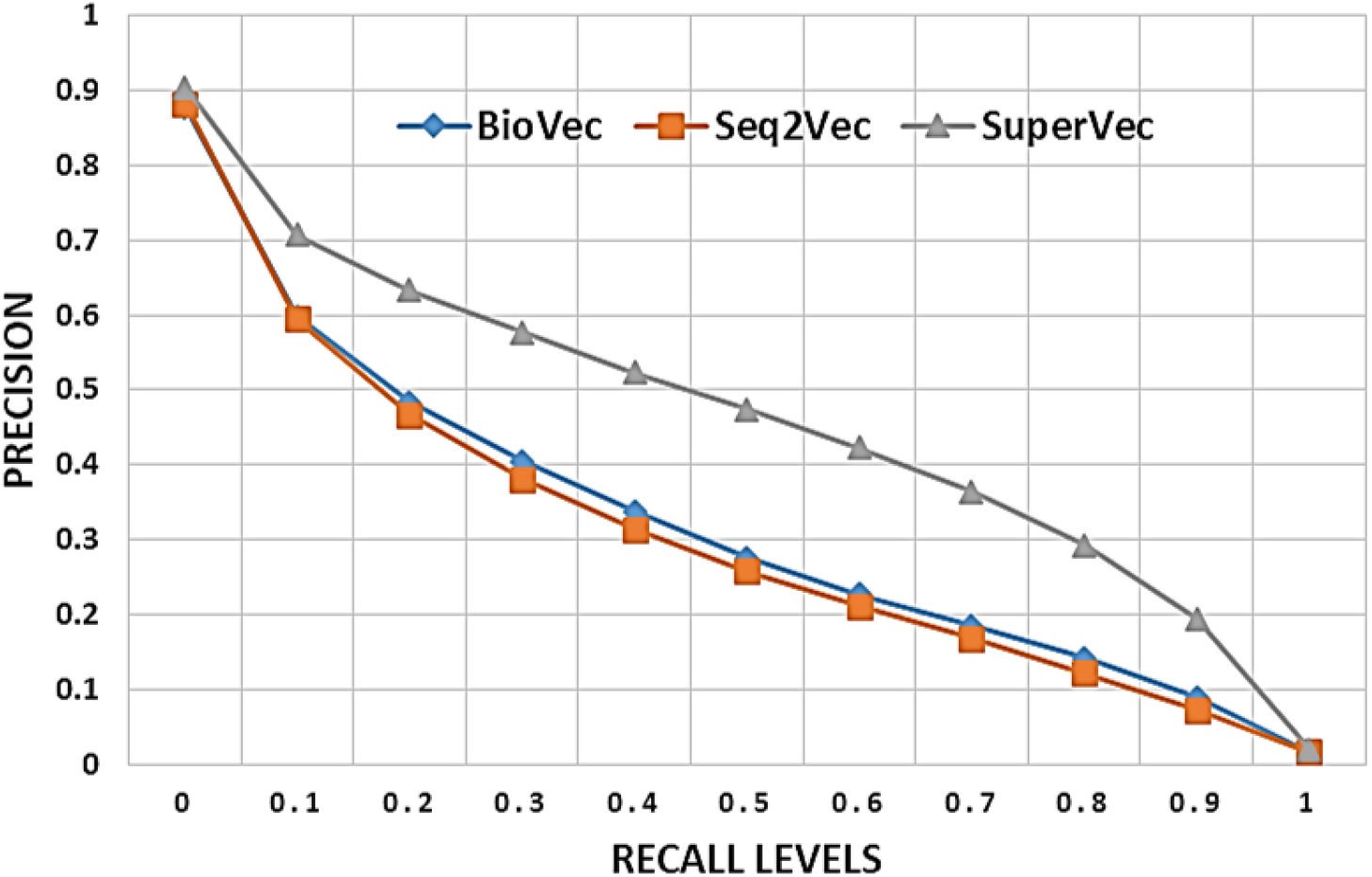
100 classes experiment: Comparison of interpolated average precision values for retrieval task performed on 100 classes database following setup 2.

### 5.2 Retrieval using H-SuperVec

`H-SuperVec` is the hierarchical version of `SuperVec` and is specifically designed to address the retrieval problem. As discussed before, `H-SuperVec` partitions the set of classes in the database into a series of exclusive and exhaustive subsets to make a binary tree (Fig 3). We then train a `SuperVec` model for each of the subset nodes of the tree. For a set of classes, many such partitions are possible, and the choice of partitioning may affect the training of `SuperVec` networks, which in turn affect the query-subject similarity computation and subsequent retrieval performance. To analyze the effect of partitions on retrieval results, we perform the retrieval experiment on a randomly chosen set of eight classes, with the experimental protocol following setup 1. We partition the chosen classes into equal-sized subsets and compute the precision-recall values using the `H-SuperVec` approach for all of the 35 possible partitions. To calculate the query-subject similarity (Eq. (7)) we follow the process explained in Fig 4 and assign the weights to node1, node2 and node3 as 0.75,0.125,0.125 respectively. We compare the precision-recall values obtained by employing `H-SuperVec` against `SuperVec` and report the percentage change in the precision value at different recall levels. The results obtained from all 35 partitions show a similar trend; Fig 9 shows the plot for five of these splits. These results demonstrate that:

- The choice of the partition at the root node has a negligible impact on the performance of `H-SuperVec`; in other words, a random partition may be chosen for applying `H-SuperVec`.

**Fig 9.**
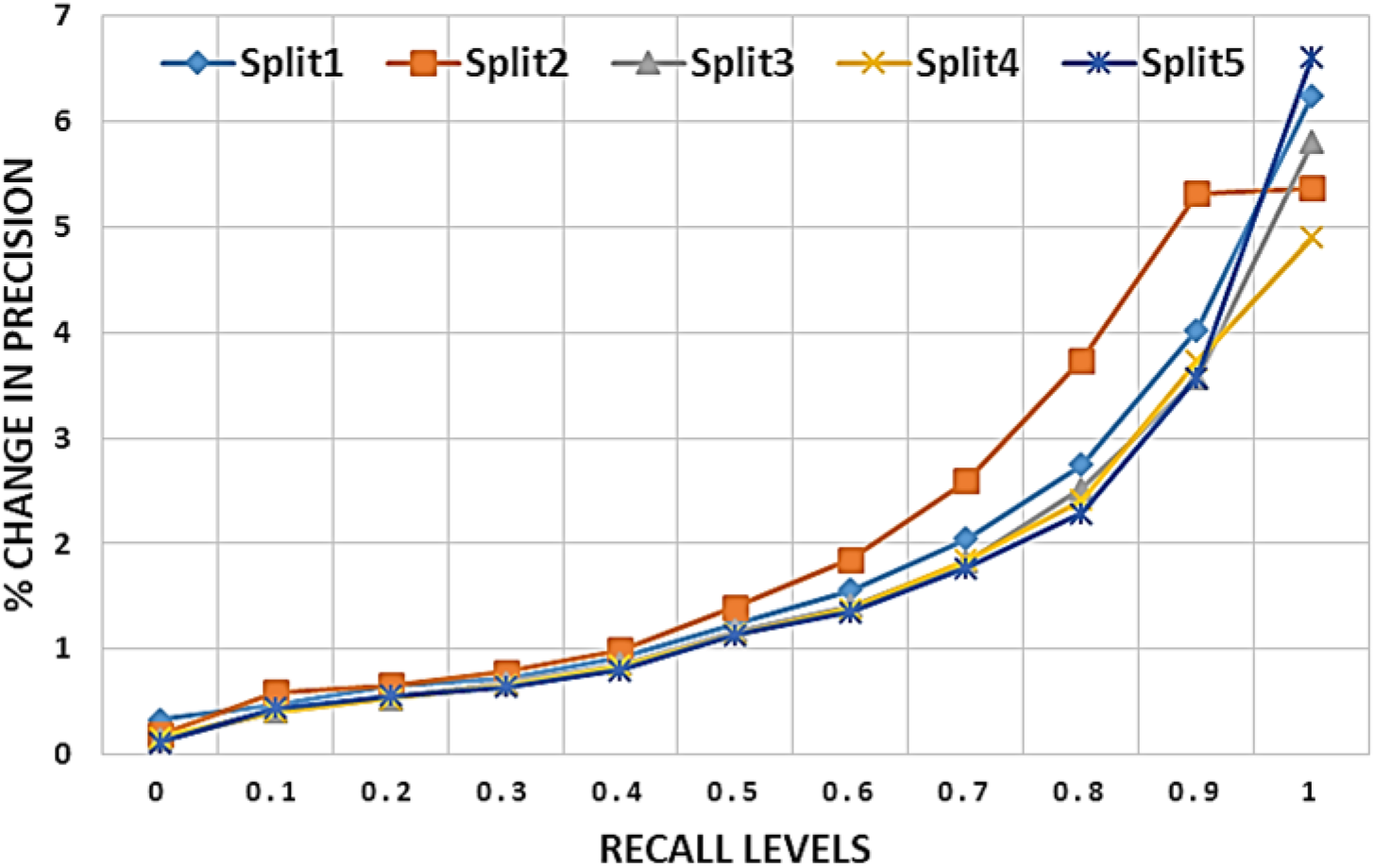
Percentage change in precision values: These plots shows the percentage change in precision values obtained for `H-SuperVec` as compared to `SuperVec` for the retrieval task performed on randomly chosen eight classes. The graph is shown for five splits among possible 35 splits.

Based on the observation made from the eight class experiments, we use a random partition of equal size for applying `H-SuperVec` on larger databases containing the largest 50, 100, 150 and 200 classes. The results for the 100 and 200 class database are shown in Fig 10. Similar to the eight class experiment, we observe that `H-SuperVec` approach outperforms `SuperVec`, giving an improvement in precision values as high as 40%. These results validate our claim that using multiple observations of the distance give a better measure of the similarity of the sequences in the embedding space leading to performance superior to `SuperVec` on the retrieval task.

**Fig 10.**
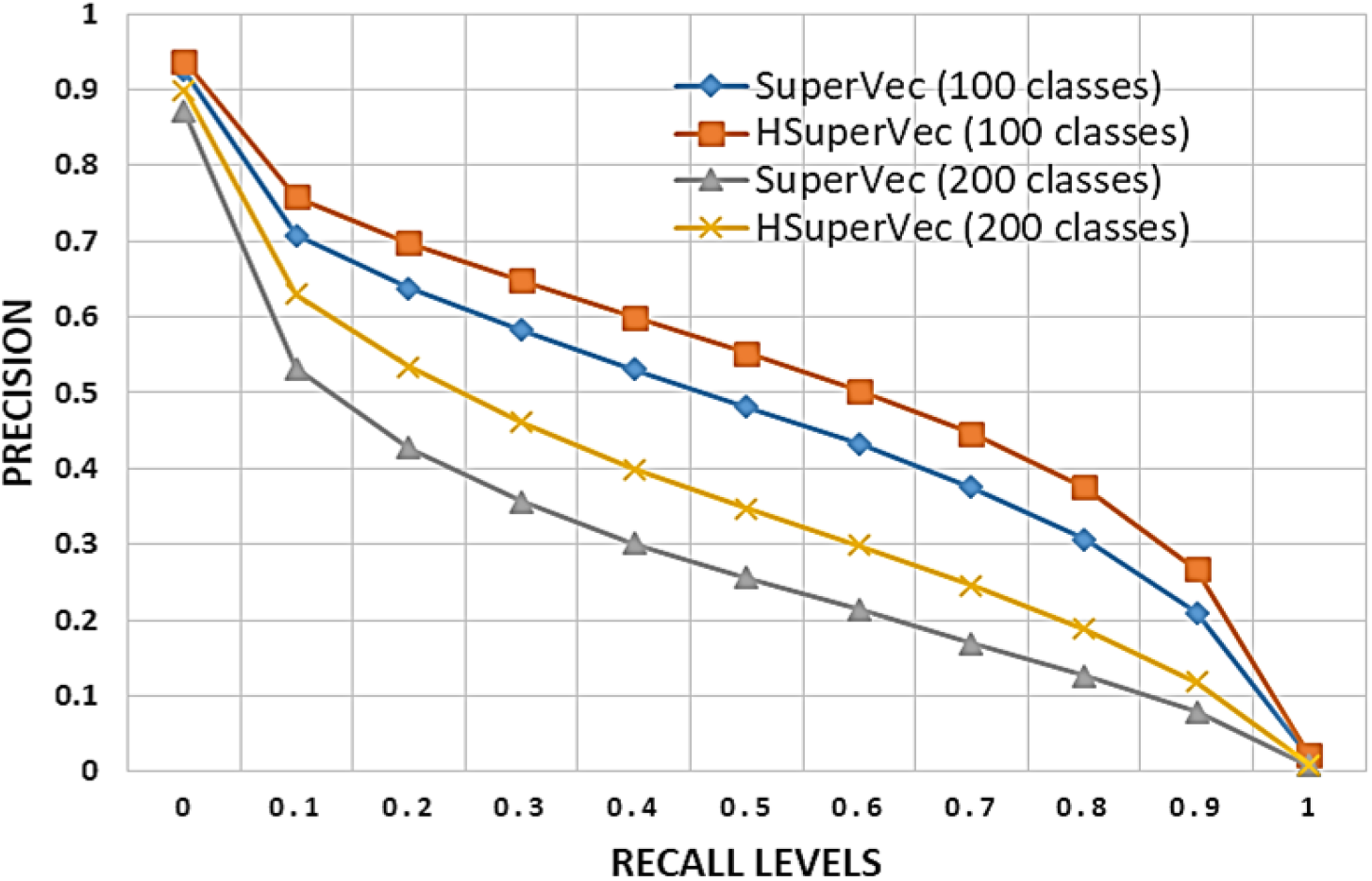
H-SuperVec vs SuperVec: Comparison of interpolated average precision values for retrieval task performed on 100 and 200 classes database.

We consider the querying time required for each of these methods in the section below.

### 5.3 Querying Time

Querying times on a retrieval task for a database of ~ 90*k* sequences and 200 classes are shown for all methods in Table 7. Timings reported are averaged over a set of 1108 randomly selected queries, chosen in equal proportion from each class. Querying times are seen to be similar across the Representation Learning methods. Since the `SuperVec` models required for `H-SuperVec` are generated in parallel, the querying time for `H-SuperVec` remains approximately the same as for `SuperVec`. Nearest neighbour search requires the same amount of processing time for all RL methods.

**Table 7.** Querying Time: Mean querying time (in milliseconds) computed for 1108 queries for the database of size 90*k* (200 classes).

| Methods | BioVec | Seq2Vec | SuperVec | H-SuperVec | BLAST |
| --- | --- | --- | --- | --- | --- |
| Querying Time | 38.30 | 20.11 | 20.11 | 22.09 | 765.19 |

When compared to BLAST, RL approaches takes ~ 35× lower time in processing a query. Note that the querying time reported in Table 7 is computed when the queries are processed serially. For vector-based methods, multiple queries can be processed together thus further reducing the querying time. For the above mentioned query set, when all queries are processed together we note that RL method took only 0.77 *sec*, i.e., the average querying time is 0.69 *msec* which thus gives us ~ 1000× speedup when compared to BLAST.

The advantages offered by `H-SuperVec`—the gain in processing time and the improved retrieval results when compared to the earlier models (`SuperVec, BioVec`, and `Seq2Vec`) —suggest that we can use it along with BLAST for significantly faster retrieval of sequences with only a modest reduction in fidelity of the results. We now consider this hybrid approach in detail.

### 5.4 Hybrid Approach: HSuperVec+BLAST

The proposed hybrid approach—`H-SuperVec+BLAST` (`H+BLAST`)—follows a two-step retrieval process. `H-SuperVec` is utilized initially to prune the original database to produce a list of results potentially relevant for the given query. Here selection is based on nearest neighbour search in the database embedding space. The list of possible relevant subjects (the reduced database) obtained via `H-SuperVec` is then provided to BLAST for re-ranking in accord with the given query. Fig. 11 shows the block diagram for the `H+BLAST` approach, where *DB* represents the original database, *DB_r_* the reduced database set obtained from the first step and *DB_o_* the final ranked list of similar sequences for the given query.

**Fig 11.**
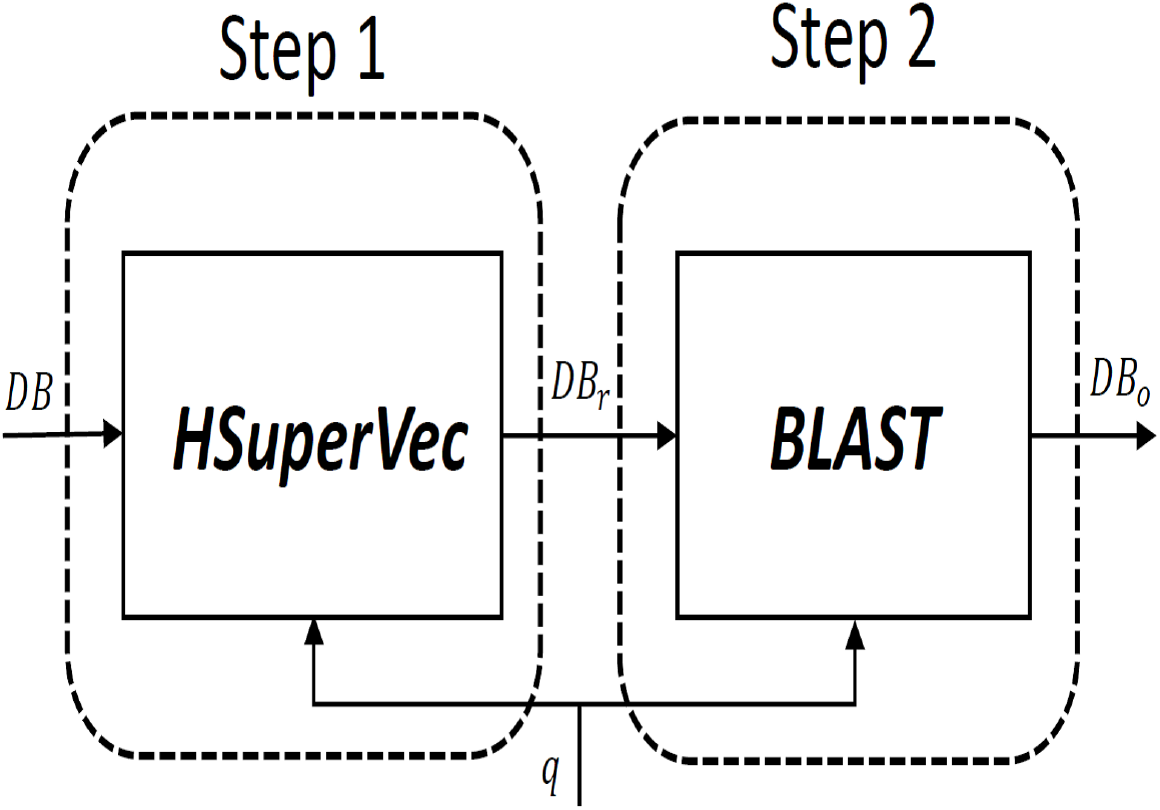
Hybrid Approach: Step1 uses `H-SuperVec` for pruning the original database (*DB*) and gives reduced database (*DB_r_*). In step2, BLAST re-ranks *DB_r_* based on alignment-based similarity between its sequences and the given query, *q* and finally provide the list of retrieved sequences, *DB_o_*.

The size of the *DB_r_* is fixed by choosing a sufficient number of nearest neighbors (NN) for a given query. For our experiment on 90*k* sequences and 200 classes, we keep NN = 10*k*, thereby allowing BLAST to operate on a comparatively small database (*DB_r_*). This provides a significant improvement in querying time compared to the direct application of BLAST to the original database. The average querying times using `H+BLAST` and BLAST to process 60*k* queries on a database of 90*k* sequences are ~ 400 *msec* and 1 *sec* respectively. Also, the use of BLAST in step 2 gives a performance improvement over `H-SuperVec` on the retrieval task. With the increase in database sizes, we expect to see further improvements in querying time and retrieval performance of `H+BLAST` as compared to `BLAST` and `H-SuperVec` respectively. Utilizing both approaches together thus gives us the best of both worlds —faster processing of queries and higher precision levels in the results.

Fig 12 provides the comparison for the methods considered of retrieval performance over the database of 200 classes. As shown, the proposed approaches, `SuperVec, HSuperVec` and `H+BLAST` considerably improve over the baseline RL approaches; Fig 13 shows the percentage gain in precision value achieved by our approaches compared to the baseline approaches.

**Fig 12.**
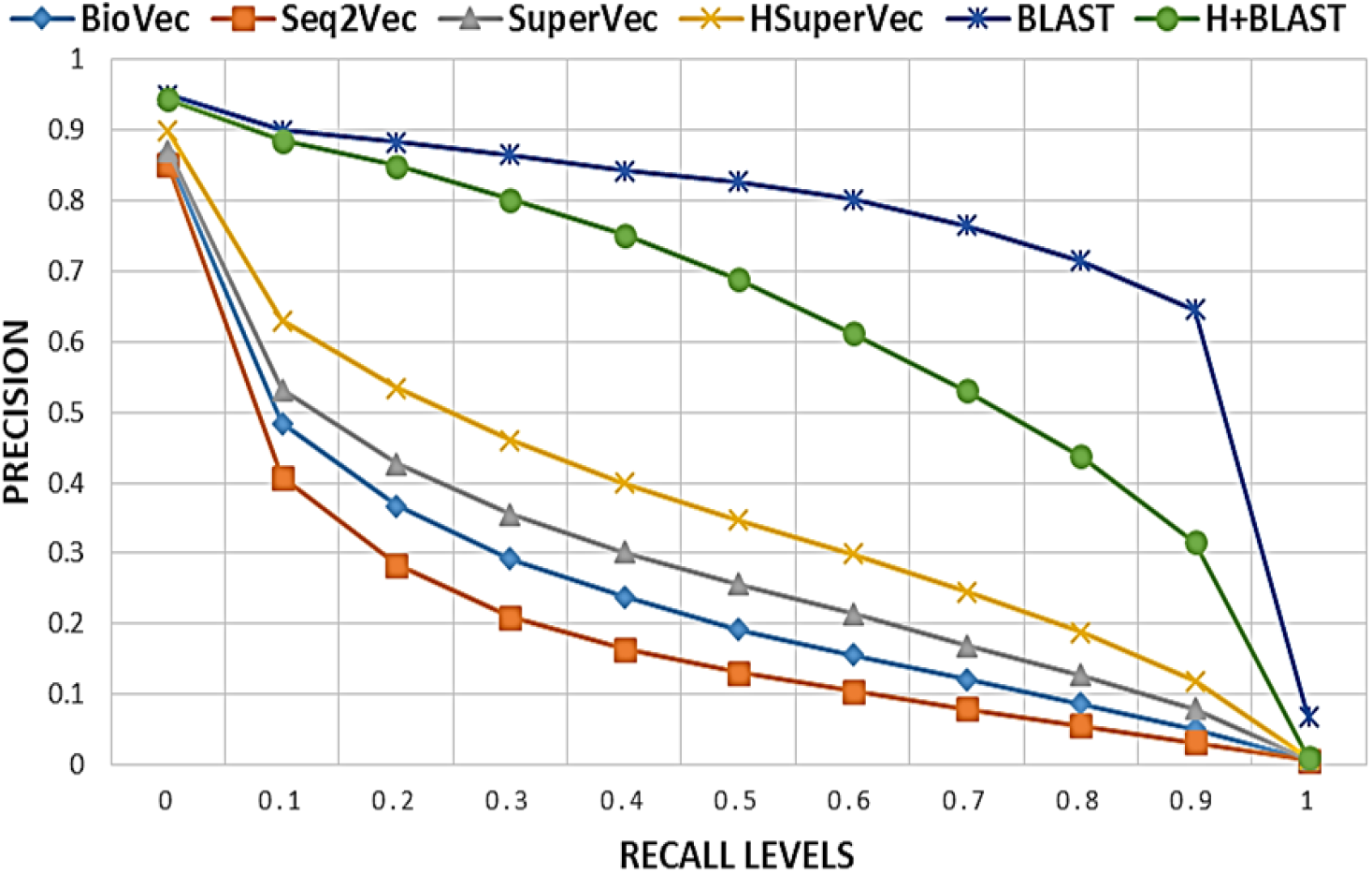
Retrieval performance comparison: The results are averaged over 60*k* queries on the database of 90*k* sequences and 200 classes.

**Fig 13.**
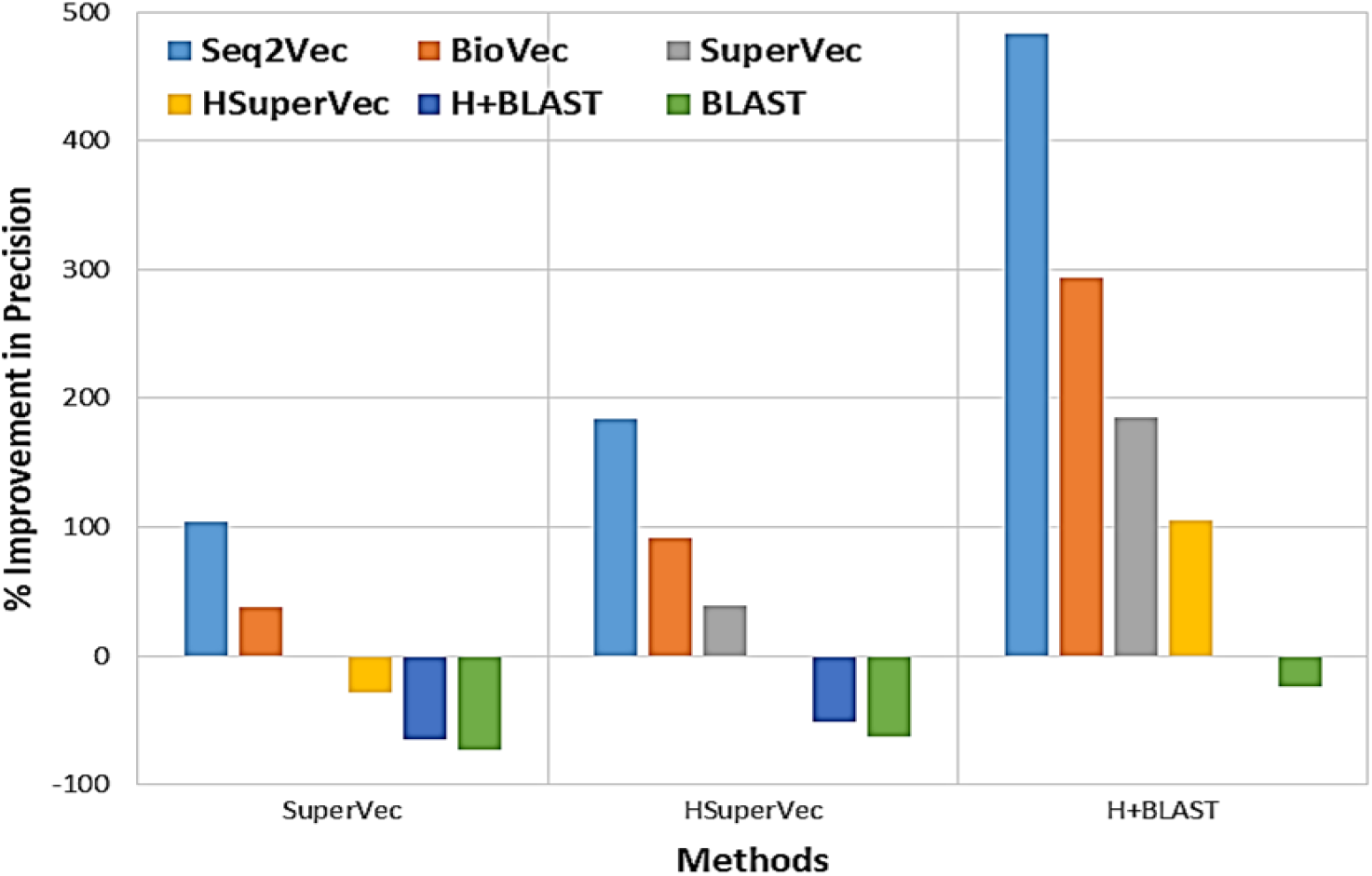
Improvement in precision: The plot shows the percentage improvement in precision value achieved by `SuperVec, HSuperVec` and `H+BLAST` over other methods. Precision is compared at 0.6 recall for the database of 90*k* sequences and 200 classes.

Improved RL approaches are expected to yield better database and query embeddings, so that the relevant subjects for a given query may be confined to a close neighborhood in the database embedding space, leading to improved performance on the retrieval task. Improvements in RL approaches would also allow us to choose a smaller number of candidate relevant subjects for a given query, thus providing further gains in processing time and performance for hybrid approaches.

## 6 Conclusions

In this paper, we introduced `SuperVec`, a supervised approach to learning embeddings for biological sequences. `SuperVec` not only learns from the information present in the sequences, but also captures sequence label information. We have demonstrated the utility of `SuperVec` generated embeddings for the homologous sequence retrieval task and noted its superior performance relative to other RL approaches in the bioinformatics domain. The better performance of `SuperVec` on the retrieval task suggests that we can generate task-specific sequence embeddings by infusing relevant meta information along with the sequence information during the learning phase of a RL model.

We also presented a hierarchical version of `SuperVec`—specifically designed for the sequence retrieval task—which we call `H-SuperVec. H-SuperVec` is seen to provide improved performance `SuperVec` when applied to the retrieval task. The major advantage offered by our methods lies in the speedy retrieval of relevant sequences from the database for a given query, some 30-50× faster than BLAST. We also proposed a hybrid approach for the retrieval task which exploits these performance advantages. `H-SuperVec` can quickly provide a list of the possible relevant subjects from a large database which can be then re-ranked with BLAST. We show that such an approach can give us early precision comparable results to BLAST along with much-improved querying speed.

Use of representation learning approaches for sequence comparison is growing, and the improvement in retrieval performance achieved with our approaches is a strong step forward in this direction. Our current work is focused on further improving the Representation Learning techniques for biological sequences so that we can obtain a better estimate of sequence similarity from the distances calculated in the embedding space. Embeddings generated through such RL techniques can then be used for a wide range of downstream bioinformatics tasks, including large scale phylogeny construction, PPI prediction, and sequence clustering.

